# Proper migration of lymphatic endothelial cells requires survival and guidance cues from arterial mural cells

**DOI:** 10.1101/2021.06.30.450504

**Authors:** Di Peng, Koji Ando, Marleen Gloger, Renae Skoczylas, Naoki Mochizuki, Christer Betsholtz, Shigetomo Fukuhara, Katarzyna Koltowska

## Abstract

The migration of lymphatic endothelial cells (LECs) is key for the development of the complex and vast lymphatic vascular network that pervades most of the tissues in an organism. In zebrafish, arterial intersegmental vessels together with chemokines have been shown to promote lymphatic cell migration from the horizontal myoseptum (HM). Here we found that LECs departure from HM coincides with the emergence of mural cells around the intersegmental arteries, raising the possibility that arterial mural cells promote LEC migration. Our live imaging and cell ablation experiments revealed that LECs migrate slower and fail to establish the lymphatic vascular network in the absence of arterial mural cells. We determined that mural cells are a source for the C-X-C motif chemokine 12 (Cxcl12a and Cxcl12b) and vascular endothelial growth factor C (Vegfc). We showed that ERK, a downstream component of Vegfc-Vegfr3 singling cascade, is activated in migrating LECs and that both chemokine and growth factor signalling is required for the robust migration. Furthermore, Vegfc-Vegfr3 has a pro-survival role in LECs during the migration. Together, the identification of mural cells a source for signals that guide LEC migration and survival will be important in the future design for rebuilding lymphatic vessels in the disease contexts.

## INTRODUCTION

The lymphatic vessel network spans across the whole body to balance the tissue fluid homeostasis, coordinate the immune responses, and enable dietary fat absorption in the intestine. The robustness of the formation and reproducibility of the vascular tree is dependent on molecular dynamics and tissue-tissue interaction required for the precision and fine-tuning of lymphatic endothelial cell (LEC) migration. Although multiple previous studies have uncovered important signals and cells guiding lymphatic vessel formation (Bussmann et al., 2010; Cha et al., 2012; Jafree et al., 2021), the recent technological developments and new transgenic lines (Ando et al., 2016; Wang et al., 2020) have opened up opportunities to identify further regulators of LEC migration.

In zebrafish trunk, lymphatic vessel specification is marked by the expression of transcription factor Prox1 in ECs around 32 hours post fertilization (hpf) in response to Vegfc-Vegfr3 signalling (Koltowska et al., 2015a). Around 34 hpf, venous-derived Prox1 positive cells sprout from the posterior cardinal vein (PCV) and migrate to the horizontal myoseptum (HM), establishing parachordal lymphatic endothelial cells (PL) (Hogan and Schulte-Merker, 2017). After about 10 hours the PLs move out from HM and migrate dorsally or ventrally and by 5 days post fertilization (dpf) give raise to the main trunk lymphatic vessels, including dorsal longitude lymphatic vessel (DLLV), intersegmental lymphatic vessel (ISLV) and thoracic duct (TD), by 5 days post fertilization (dpf) (Kuchler et al., 2006; Yaniv et al., 2006). During this later migration, vast majority of the LECs are associated with the arterial intersegmental vessels (aISV) (Bussmann *et al*., 2010). In mutant embryos lacking aISV (*plcy*^*t26480*^ and *kdrl*^*hu5088*^ mutants), PLs remain in HM, and subsequent the lymphatic network formation is compromised (Bussmann *et al*., 2010). On molecular level, Cxcl12b secreted from arterial ECs (aECs) in ISV guides this LECs migration via Cxcr4 receptor expressed in LECs (Cha *et al*., 2012). Yet, if other tissues or cells cooperate with arterial ECs to support this migration remains unknown.

Simultaneous with LEC development described above, vascular mural cells (MC) are formed *de novo* along aISVs and beneath the dorsal aorta. aISVs play a critical role in this process, and MC-emergence is completely abolished in the absence of arterial ECs (Ando *et al*., 2016). The spatio-temporal similarity of MC and lymphatic vessel development around aISV raises the question about possible interaction between MCs and LECs in this region.

VEGFC-mediated signalling through the VEGFR3 receptor is essential for multiple steps of lymphatic vessels formation, including LEC proliferation, differentiation, and migration. *In vitro* cell culture experiments have demonstrated that VEGFC-VEGFR3 and β1 integrin to promote LEC migration (Makinen et al., 2001; Wang et al., 2001). Studies using *vegfc* reporter zebrafish line has uncovered multiple sources of *vegfc*, including the fibroblasts and neurons, which contributes to the initial sprouting and migration of lymphatic vessel into the HM (early migration) (Wang *et al*., 2020). The requirement of VEGFC-VEGFR3 and its source(s) in the LEC migration out of the HM (late migration) remains to be defined. Mechanistically, transcription factor Mafba, which regulates LEC migration but not proliferation, has been shown to act downstream of VEGFC-VEGFR3 signalling (Dieterich et al., 2015; Koltowska et al., 2015b). A genome-wide analysis further indicted the presence of a transcriptional network controlling LEC migration, through the induction of chemokine receptors that promote chemotaxis in migrating LECs (Williams et al., 2017). Although the migratory regulators have been identified, the upstream cellular source of the signals initiating the migration is unknown.

Here, we took advantage of the transgenic zebrafish reporters which allow to visualize MC and LEC simultaneously at high spatio-temporal resolution *in vivo*, and to investigate their communication during lymphangiogenesis. We found that MCs emergence precedes LEC migration along aISV and that LECs interact with MCs residing at the aISVs. Moreover, in the absence of MCs, the LECs migration was inhibited, and lymphatic vessel formation was compromised. We further determined that MCs produce lymphangiogenic factors including *vegfc, cxcl12a*, and *cxcl12b*. Thus, this study uncovers a close interaction between MC-LEC of functional importance for lymphatic vessel formation in the zebrafish trunk.

## RESULTS AND DISUCSSION

### MCs and LECs interact during LEC migration

To address a potential interaction between MCs and LECs around arteries, we examined their distribution around aISVs, using the reporter lines *Tg(lyve1b:DsRed);Tg(flt1:YFP);Tg(pdgfrb:GFP)* and found the spatial proximity, with MCs being sandwiched between the aISV and the migrating LEC at 4 dpf (**Figure 1A**). Time lapse imaging showed that LECs migrated out from HM immediately after the emergence of *pdgfrb*^+^ MCs (**Figure 1B and C, Supplemental Figure 1A**). The number of MCs was not changed before and after the LEC migration along aISV (**Figure 1D**) and in approximately 90% of the cases, LEC migrated towards and interacted with the MC residing on aISV (n=24 **Figure 1E-F**). When LEC migrated out from HM, we noticed that LEC dynamically extended and regressed protrusions as if they sensed the environment and searched for the nearby located MC (**Figure 1E, Supplemental Figure 1A**). To understand the biological significance of this interaction, we quantified the velocity of LEC migration along aISV and found that the LECs in contact with the MCs migrated two times faster than LECs migrating along aISV without MCs (**Figure 1G**). These observations suggest that MCs might provide directional cues to promote robust LEC migration.

**Figure 1.**
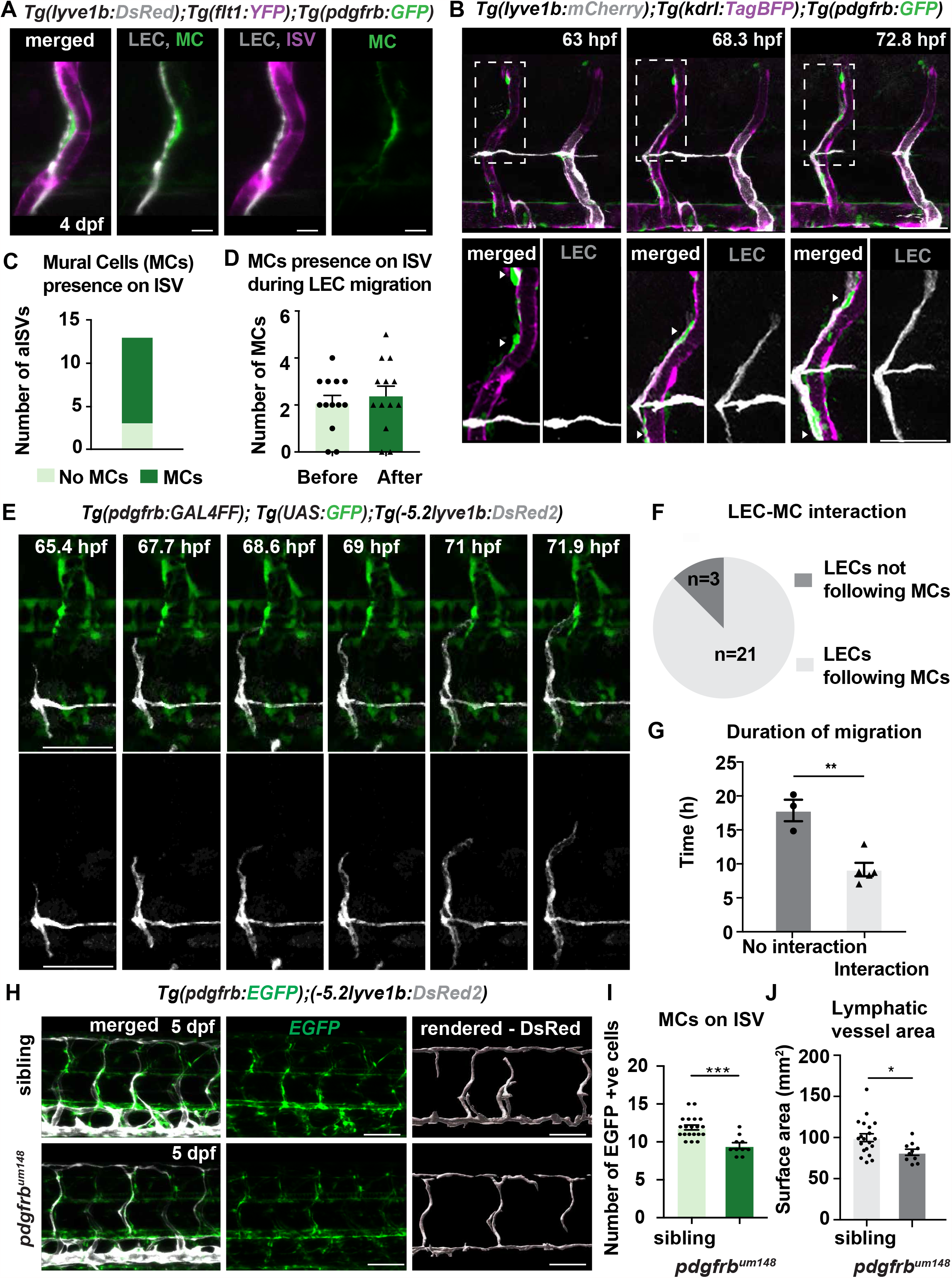
*pdgfrb* ^high^ mural cells emerge around aISVs prior to LEC migration and provide guidance. (A) Confocal stack image of trunk aISV in 4 dpf *Tg(−5*.*2lyve1b:DsRed);Tg(flt1:YFP); TgBAC(pdgfrb:GFP)* of lymphatic endothelial cells (gray, LEC), arterial intersegments vessels (magenta, aISV) and MCs (green, MC). Scale bar; 10 μm. (B) Confocal stack images from time-lapse images in the trunk of 2 dpf *Tg(lyve1b: mCherry);Tg(kdrl:TagBFP);TgBAC(pdgfrb:GFP)* embryo. Boxed regions are enlarged in the bottom. Arrowheads indicate *pdgfrb*^+^ MCs (green) around aISV (magenta) prior to (bottom left) and during (bottom middle and right) LEC (gray) migration. Scale bars; 50 μm or 30 μm (enlarged image). (C) Quantification of aISV with (n=10) or without (n=3) MCs presence when LECs left HM for time lapse videos as in (E). (D) Quantification of MC number around aISV (n=13) at the start and end of the migration for time lapse videos as in (E). (E) Confocal stack images from time lapse of LEC migration. *TgBAC(pdgfrb:GAL4FF);(UAS:GFP)* in green and *Tg(−5*.*2lyve1b:DsRed2)* in grey. Scale bar; 50 μm. (F) Quantification of LEC and MC interaction during migration. Migrating following MC n=21, migrating not following MC n=3. (G) Quantification of duration of LEC migration with (n=5) or without (n=3) contacting MCs. Data are presented as mean ± SEM. **p<0.005 (H) Confocal stack images of *Tg(pdgfrb:GAL4FF); Tg(UAS:GFP)* (green) and *Tg(−5*.*2lyve1b:DsRed2)* (grey) in the trunk of sibling (top) and *pdgfrb*^UM148^ mutant (bottom) embryos at 5 dpf. Lymphatic structure is rendered with *lyve* channel in IMARIS (right panel). Scale bar; 100 μm. (I) Quantification of number of *pdgfrb*^+^ mural cells around ISVs in siblings (n=20) and *pdgfrb*^UM148^ mutant (n=10). Data are presented as mean ± SEM, unpaired two-tailed Student’s t-test was used. ***p<0.0001. (J) Quantification of surface area of lymphatic vasculature in siblings (n=20) and *pdgfrb*^UM148^ mutant (n=10). Data are presented as mean ± SEM, unpaired two-tailed Student’s t-test was used. *p<0.05.

### MCs promote lymphatic vessel formation

We next asked if *pdgfrb* positive MCs are necessary for lymphatic vessels formation. PDGFRβ is known to be essential for MC development, especially their proliferation and migration (Ando *et al*., 2016; Gaengel et al., 2009). The *pdgfrb*^*um148*^ mutant zebrafish (Kok et al., 2015) showed 30% reduction of MC-number around aISVs (**Figure 1H-I)**. Coincidently, trunk lymphatic vasculature formation in *pdgfrb*^*um148*^ mutant zebrafish revealed slight reduction in the network formation. The rendering of the area of *lyve:DsRed* labelled lymphatic vessels, as a measure of lymphatic vessels density, revealed on average an area of 100 mm^2^ in the sibling vs. 70 mm^2^ in the *pdgfrb*^*um148*^ mutants (**Figure 1H and 1J**). Treatment with a PDGFRβ inhibitor, AG1296 from 48 hpf, led to a greater reduction in MC-coverage and LECs number (**Supplementary Figure 1B**). Together, although it cannot be excluded that AG1296 inhibitor directly affected lymphatic vessels development, these observations suggest a requirement of *pdgfrb*^*+*^ MCs for lymphatic development.

As both mutant or AG1296-treated larvae retained a substantial proportion of their MCs, we decided to eliminate *pdgfrb*^+^ MCs utilizing MC-selective Nitroreductaes (NTR) and metorodinazole (Mtz) system (NTR-system) (Curado et al., 2008), *TgBAC(pdgfrb:Gal4FF);Tg(14xUAS:3xFLAG-NTR, NLS-mCherry)* to confirm the involvement of MCs for lymphatic vessel formation. In this transgenic line, Mtz is converted to its cytotoxic form by NTR expressed in *pdgfrb*^+^ MCs leading to selective MC-death in the trunk (**Supplementary Figure 2A**). When ablating MCs just prior to LEC migration out from HM by utilizing this MC-selective NTR-MTZ ablation system, LEC migration along aISV and subsequent TD formation were severely compromised (**Figure 2 A-C**). To further determine if MCs are necessary for LEC migration, we ablated MCs locally by multi-photon laser after LEC just migrated out of HM (**Figure 2D**). As a control we targeted the tissue adjacent to the MCs in the same embryo (**Figure 2E**). We imaged the same embryos one day later and observed that 40% of the LECs in the MC-ablated group fail to migrate to form DLLV (**Figure 2E-H**). The time-lapse imaging over 5 hours post ablation (pa) confirmed that LEC migration was dramatically inhibited in the MC-ablated group compared to the control non-MC ablated group (**Figure 2E-H**).

**Figure 2.**
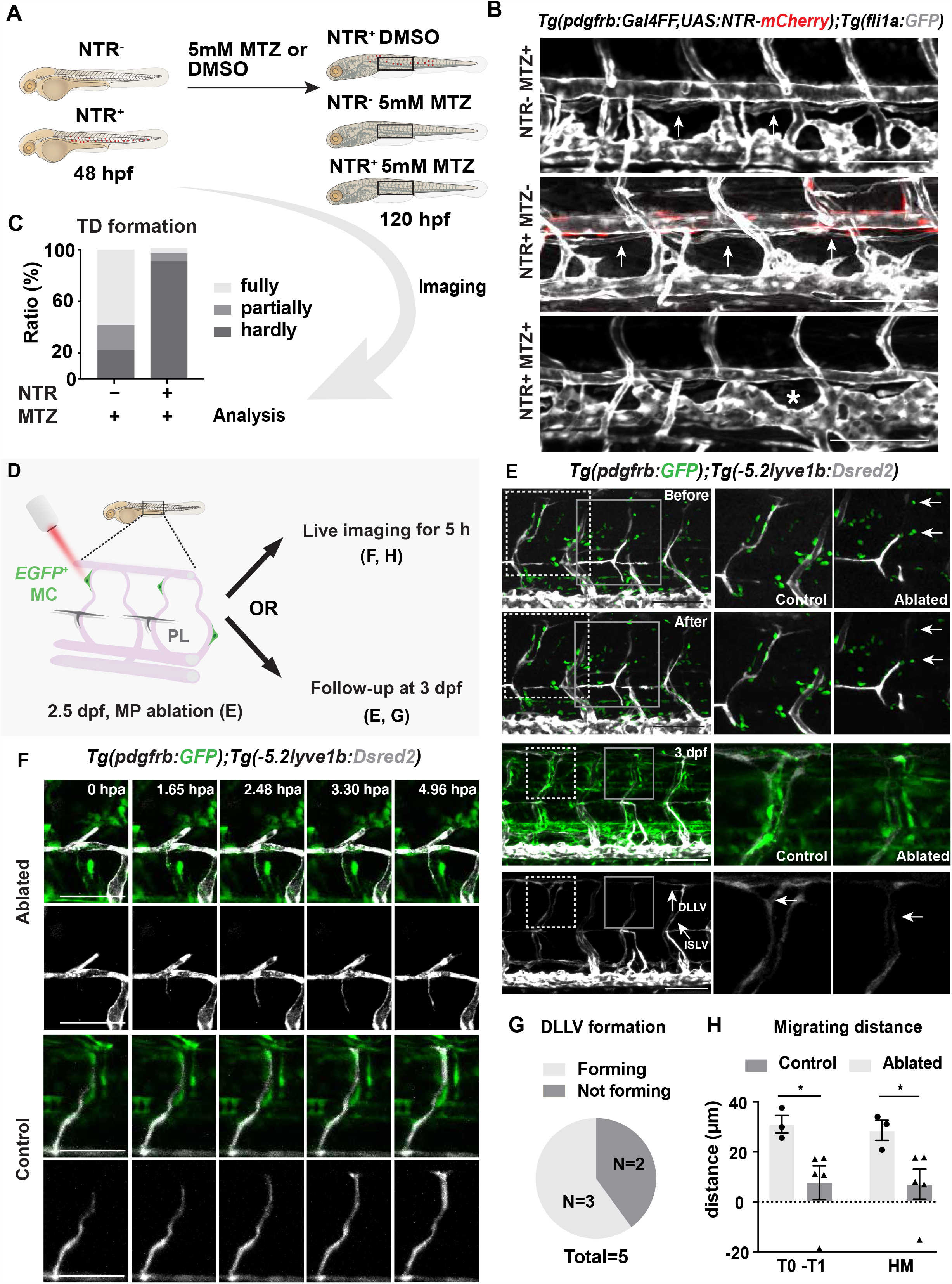
Mural cells are required for robust formation of lymphatic vascular bed. (A) Work flow of cell ablation by NTR-MTZ system. *Tg(pdgfrb:Gal4FF);Tg(14xUAS:3xFLAG-NTR,NLS-mCherry)* (red) and *Tg(fli1a:GFP)* (grey) were imaged at 120 hpf after treatment of DMSO or 5mM MTZ from 48 hpf, and the formation of thoracic duct (TD) was analyzed. (B) Confocal stack images of the trunk in 5 dpf embryos treated as described in (A). Arrows indicates TD forming beneath dorsal aorta. Asterisks indicates the absence of TD. Scale bar; 100 μm. (C) Quantification of (B). Embryos were scored as fully, partially and hardly formed based on the TD development. Data were presented as ratio to total number of embryos analyzed. (D) Work flow of cell ablation by multi-photon microscopy. Mural cells (MC, green) labelled by *TgBAC(pdgfrb:GAL4FF);(UAS:GFP)* and PL by *-5*.*2lyve1b:*DsRed2 (grey) embryo. The MC on intersegmental vessels in proximity to PL were ablated at 57 hpf. For analysis, ablation was either followed by time-lapse imaging or confocal imaging at 3 dpf. (E) Confocal stack images of before and after ablation (top two raw) and follow-up at 3 dpf (bottom two row). Control ablation (dashed box) at the adjacent region of GFP^+^ MCs and GFP+ MC on aISV ablation (solid grey box) was performed in the same embryos. Arrowheads indicate ablated GFP-positive cells and development of targeted LEC at 3 dpf. Scale bar; 100 μm. Middle and right panel, zoom-in images cropped in z-stacked. (F) Live imaging of the lymphatic endothelial cells (LEC) migration in the context of control (top images) and GFP+ MC on aISV (bottom images) ablation, confocal stack images from time lapse at selected timepoints from 0 hours post ablation (hpa) to 4.96 (hpa). Scale bar; 50 μm. (G) Quantification of ablated LEC forming DLLV at 3 dpf. DLLV forming (n=3), not forming (n=2). (H) Quantification of migrating distance from time-lapse videos corresponding in (D). Distance was calculated as both T0-T1 and the perpendicular distance between the T1 and HM for embryos with (ablated, n=4) or without (control, n=3) ablation. T0, the sprouting front of LEC at the start of video; T1, sprouting front of LEC at the end of video. Data are presented as mean ± SEM, unpaired two-tailed Student’s t-test was used on two types of measurements respectively. *p<0.05.

Together, these data show the MC-LEC interaction at high spatio-temporal resolution during lymphatic development and demonstrate an important role of arterial associated MCs for the robust LEC migration.

### Chemokines are expressed in *pdgfrb*^+^ mural cells during LEC migration

The importance of MCs for LEC migration might represent a direct interaction between the two cells or an indirect effect mediated via aISV ECs, which were previously shown to be necessary for LEC guidance (Bussmann *et al*., 2010). The absence of MCs does not affect arterial identity at early stages in zebrafish (Ando et al., 2019), arguing that the importance of MC in LEC development is not simply to regulate aEC presence or abundance. However, in order to directly test aEC function is critical for MC-dependent LEC migration in aISV, we also ablated ECs in aISV after the emergence of MCs using the multi-photon laser system. We observed that even in the absence of aEC, but remaining presence of MCs, LEC migration progressed (**Figure 3A**), suggesting that signals from MCs are sufficient to promote LEC migration. Thus, the previously reported strong inhibition of LEC development in aISV-depleted mutants (Bussmann *et al*., 2010) might include the effects of MC-loss as aISVs ECs are essential for MC-formation (Ando *et al*., 2016). However, as the aISV rapidly regrow following laser ablation system (**Figure 3A**-**(C**), the long-term effects cannot be assessed, and it remains to be determined to what extend the molecular signals from MC act in synergy with aECs to promote LEC migration.

**Figure 3.**
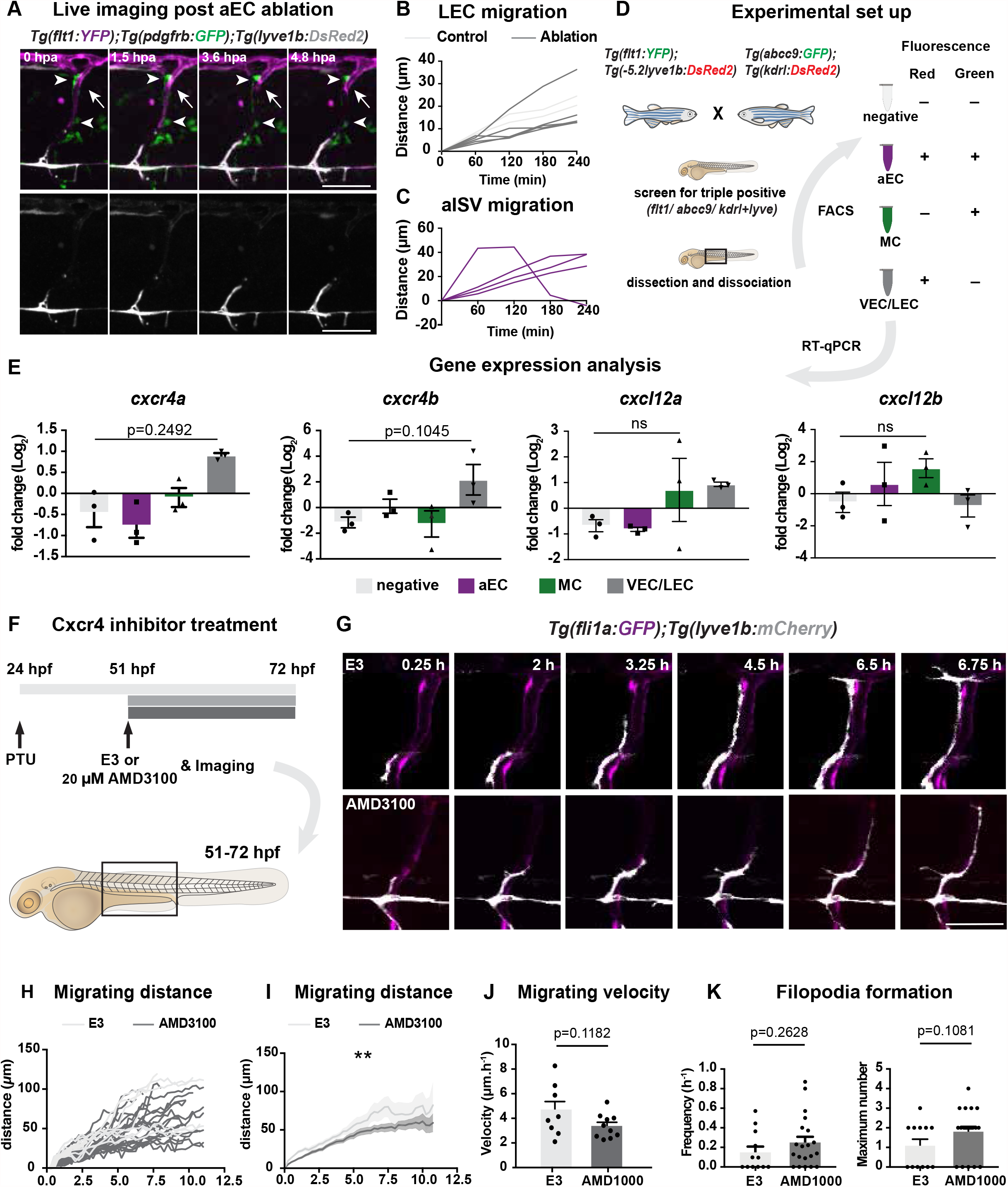
Chemokines are expressed in ISV-mural cells during LEC migration. (A) Confocal stack images from time lapse post multi-photon laser ablation in 2 dpf *Tg(flt1:YFP)* (magenta); *TgBAC(pdgfrb:GFP)* (green) and *Tg(−5*.*2lyve1b:DsRed2)* (grey). Arrowheads indicate remained GFP^+^ mural cells without aISV. White arrows indicate the ablated site of aISV. Scale bar; 50 μm. (B-C) Quantification of LEC (n=4) and aISV (n=4) migration distance post two-photon laser ablation. (D) Illustration of FACS sorting and qPCR analysis on 3 dpf embryos. (E) qRT-qPCR of *cxcr4a, cxcr4b, cxcl12a* and *cxcl12b* in FACS sorted population cells at 3 dpf. Graph represents gene expression relative to geometric average of *rpl13* and *β-actin* from three biological repeats (mean ± SEM). Kruskal-Wallis test with Dunn’s post hoc test was used. No significance (ns), p≥ 0.9999. (F) Work flow of Cxcr4 inhibitor treatment. Embryos of *Tg(fli1:GFP);Tg(lyve1b:mCherry)* was grown in PTU from 24 hpf, then changed to 20 μM ADM3100 or E3 water at 51 hpf. Time-lapse imaging was started then in a two-well slide that allows spontaneous imaging for both groups. (G) Confocal stack images from time lapse imaging indicated in (F). Scale bar; 50 μm. (H) Quantification of dorso-ventral migration showing individual tracks for sprouting front of PLs in E3 water (n=10) and AMD3100 (n=25) treated embryos as described in (G). (I) Quantification of dorso-ventral migration showing average (mean) of tracks in DMSO and AMD3100 treated group. Data are presented as mean ± SEM, unpaired two-tailed Student’s t-test was used. **p<0.005. (J) Quantification of velocity of dorso-ventral migration from time lapse video described in (3F). Nuclei of PLs in E3 water (n=10) and AMD3100 (n=25) treated embryos were tracked and the distance between begin and end position of nuclei was measured then divided by duration. Data are presented as mean ± SEM. unpaired two-tailed Student’s t-test was used. (K) Quantification of frequency of filopodia formation from time lapse video described in (3F). (left) Number of protrusions other than the sprouting front was counted and divided by the corresponding duration in E3 (n=12) and AMD3100 (n=20) treated embryos. Data are presented as mean ± SEM. unpaired two-tailed Student’s t-test was used. (right) Number of protrusions other than the sprouting front was counted and divided by the corresponding duration in E3 (n=12) and AMD3100 (n=20) treated embryos. Data are presented as mean ± SEM. unpaired two-tailed Student’s t-test was used.

Based on the above observations, we hypothesise that MC may also secrete chemoattractant molecules to guide the LECs (**Figure 3A**). It has been shown that the LEC migration is driven by the chemokines Cxcl12a and Cxcl12b (Cha *et al*., 2012), which are expressed in ECs. To test if Cxcl12a and Cxcl12b are also expressed in MCs we collected cells by fluorescent activated cell sorting (FACS) from *Tg(flt1:YFP); TgBAC(abcc9:Gal4FF); Tg(UAS:GFP); Tg(lyve1b:DsRED); Tg(kdrl:mCherry)* larvae at 3 dpf and analysed candidate genes expression in aECs (double positive for green (YFP) and red (mCherry) fluorescence), MCs (single positive for green (GFP) fluorescence), and venous ECs and LECs (single positive for red (DsRED/mCherry) fluorescence) (**Figure 3D, Supplementary Figure 3A**). Although *abcc9* reporter is highly selective for ISV-MCs in the trunk, posterior notochord is also labelled (Ando *et al*., 2019). Therefore, to avoid the possible contamination of other cell types than *abcc9* positive MCs, we used micro-dissection to isolate the trunk region containing *abcc9*-positive ISV-MCs for FACS. We confirmed the identity of sorted MCs by expression of *abcc9* and *pdgfrb* (**Supplementary Figure 3B**). In line with published data, that *cxcr4a* and *cxcr4b* are expressed by LECs and *cxcl12b* by aECs (Cha *et al*., 2012). In addition, we found that *cxcl12a* and *cxcl12b* are expressed in the *abcc9-*positive MC population (**Figure 3E**). To further understand if signalling by these ligands is essential for LEC migration, we have treated embryos with a Cxcr4 inhibitor, AMD3100 (**Figure 3F, Supplementary Figure 3C**), and found LECs migrated less and their migration velocity is decreased (**Figure 3G-J**). This coincided with an increased number of filopodia formation on LEC (**Figure 3 K**), indicating loss of guidance (Meyen et al., 2015). Together our data show that MCs produce the LEC chemo-attractants driving LEC migration.

### MC and arterial derived *vegfc* promotes LEC migration and survival

Vegfc-Vegfr3 initiates a signalling cascade driving lymphangiogenesis in zebrafish and is required for sprouting and LEC migration towards the HM (Hogan et al., 2009a; Yaniv *et al*., 2006). As Vegfc has been shown to be expressed in vascular smooth muscle cells (VSMCs) in mice (Antila et al., 2017), we hypothesised that it might be expressed also in zebrafish MC. Firstly, we checked *vegfc* expression in FAC-sorted cells by RT-qPCR and found that *vegfc* expression in aISV-MCs as well as aECs (**Figure 4A**). We also found that phospho(p)-ERK staining of LEC, which is known to be activated downstream of Vegfc-Vegfr3 or Cxcl12-Cxcr4 (Spinosa et al., 2019; Xing et al., 2017), during LEC migration along aISV (**Figure 4B**). To assess if ERK activation is required for the LEC migration from HM we have used the MEK inhibitor, SL327 (**Supplementary Figure 4A**). We treated embryos at 51 hpf just prior to their migration and assessed the phenotypes by time-lapse imaging (**Figure 4C-D**). We traced the migration distance and observed a reduced number of migrating cells and increased number of cells that stalled or regressed their migration in SL327-treated embryos (n=5) (**Figure 4E**). In addition, we found a dramatic decrease of LEC cell division from 25% controls to 1.5 % precent in SL327-treated embryos (**Figure 4G**), which is in agreement with the known necessary role of Vegfc-Vegfr3 in cell proliferation (Cao et al., 1998). SL327-treatment induced cell death in 7 out of 12 cells, which was further confirmed by TUNEL staining (**Figure 4 H-I**), suggesting that during this lymphatic developmental window ERK activation acts as a LEC pro-survival factor. These effects were not observed in AMD3100 treated embryos, indicating that ERK activation may be induced mainly via Vegfc-Vegfr3 signalling rather than Cxcl12-Cxcr4 signalling. Together, these results indicate that in addition to LEC specification and sprouting from the PCV (Karkkainen et al., 2004; Koltowska *et al*., 2015a), Vegfc-Vegfr3 signalling also instructs LEC in the subsequent migratory events to establish the lymphatic vessel network in the trunk.

**Figure 4.**
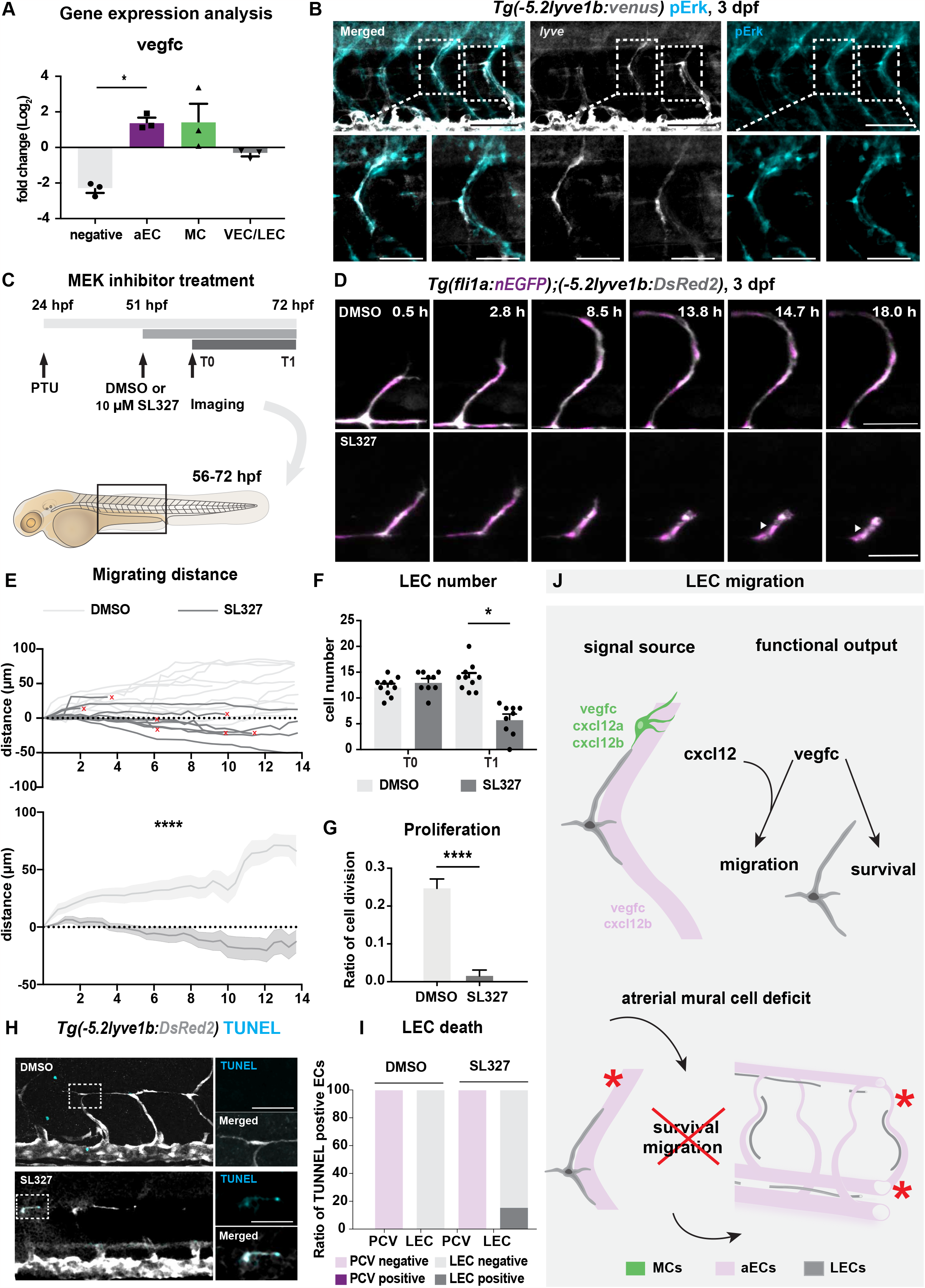
Mural cell and arterial derived *vegfc* promotes LEC migration. (A) qRT-qPCR of *vegfc* expression in trunk ECs and MCs cells at 3 dpf as described in (3E). Graph represents gene expression relative to geometric average of *rpl13* and *β-actin* from three biological repeats (mean ± SEM). Kruskal-Wallis test with Dunn’s post hoc test was used. *p<0.05 (B) Endogenous pErk (cyan, right) in migrating LEC in trunk of *Tg(−5*.*2lyve1b:venus)* embryos (α-GFP, grey, middle) at 3 dpf. Scale bar 100 μm; 50 μm in enlarged images. (C) Work flow of vegfc inhibitor treatment. Embryos of *Tg(fli1:GFP);Tg(lyve1b:mCherry)* was grown in PTU from 24 hpf, then changed to 10 μM Sl327 or DMSO at 51 hpf. Time-lapse imaging was started at 57 hpf. (D) Confocal stack images from time lapse of 57 hpf *Tg(fli1a:nEGFP)*^*y7*^ (green) and *Tg(−5*.*2lyve1b:DsRed2)* (grey) treated with DMSO or 10 μM SL327 from 51 hpf. Grey arrowheads indicate cell death. Scale bar in 50 μm. (E) Quantification of dorso-ventral migration showing individual cell tracks for nuclei of PLs (top panel) in DMSO (n=11) and SL327 (n=12) treated embryos as described in (4C). Red cross indicates cell death at the end of tracking. Bottom panel, average (mean) of tracks in DMSO and SL327 treated group. Data are presented as mean ± SEM, unpaired two-tailed Student’s t-test was used. ****p<0.0001 (F) Quantification of cell proliferation in DMSO (n=14) and SL327 treated (n=17) embryos as described in (4C). Nuclei marker in green was used to count the cell division event. (G) Quantification of total LEC number at beginning (T0) and end (T1) of the time lapse in DMSO (n=10) and SL327 (n=9) treated embryos as described in (C), data are presented as mean ± SEM. T0 DMSO vs T1 SL327 p<0.0001, T0 SL327 vs T1 SL327, T1 DMSO vs T1 SL327 p<0.0001. Other comparisons were ns, p≥0.9999. One-way ANOVA with Tukey’s post hoc test for statistical analysis. ****p<0.0001 (H) Confocal stack image of embryos treated with DMSO or SL327 as described in (4C). Embryos were fixed at 3 dpf and used for TUNEL (cyan) and α-DsRed staining (grey). Zoom-in single z stack images of TUNEL staining in merged and TUNEL channel showing the colocalization of the signal. Scale bar; 100 μm, 30 μm (enlarged images). (I) Corresponding quantification of TUNEL signal from (I). In SL327 treated group (n=45) LEC showed a positive TUNEL staining rate of 15.56% while it was 0% in DMSO treated group (n=45). (J) A model of LEC migration in zebrafish trunk. Signals promoting cell survival and migration expressed in MCs include *vegfc, cxcl12a* and *cxcl12b*at 3 dpf (top). In absence of the arterial mural cells, LECs fail to establish a robust lymphatic network at 5 dpf (bottom panel).

In summary, we demonstrated MC-LEC interaction at high spatio-temporal resolution during lymphatic development. We further uncovered an important role for artery-associated MCs in guidance of LECs, which is mediated by their secretion of chemoattractants including Cxcl12 and Vegfc. Since other sources of Cxcl12 and Vegfc have already been demonstrated in the zebrafish embryonic trunk (Cha *et al*., 2012; Wang *et al*., 2020), we propose that colonisation of the aISV by MC may provide a signalling threshold needed for LECs to depart from the HM and begin migrating along the aISVs. Our study underscores the importance of spatial and temporal control of the guidance cues and mitogens in order to promote and refine the migratory path and survival of LECs. Our finding of MC as molecular source for lymphangiogenic factors should have relevance to future designs aiming at re-establishing lymphatic vessels in disease contexts.

## Material and methods

### Zebrafish

Zebrafish were maintained in Genome Engineering Zebrafish National Facility, Uppsala University using standard husbandry conditions (Alestrom et al., 2020). Animal experiments were carried out under ethical approval from the Swedish Board of Agriculture (5.2.18-7558/14). Previously published transgenic lines used were *Tg(fli1a:nEGFP)*^*y7*^, *Tg(−5*.*2lyve1b:DsRed2)*^*nz101*^ (Okuda et al., 2012), *Tg(1xUAS:GFP)* (Asakawa et al., 2008), *TgBAC(pdgfrb:Gal4FF)*, (Ando et al., 2016) *TgBAC(pdgfrb:GFP)* (Ando et al., 2016), *TgBAC(abcc9:GAL4FF)* (Ando et al., 2019), *Tg(flt1:YFP)*^*hu4881*^ (Hogan et al., 2009b), *Tg(fli1a:GFP)*^*y1*^ (Nathan and Weinstein, 2002), *Tg(−7kdrl:DsRed2)*^*pd27*^ (Kikuchi et al., 2011), *pdgfrb*^*um148*^ (Kok *et al*., 2015), *Tg(kdrl:TagBFP)*^*mu293Tg*^ (Matsuoka et al., 2016), *Tg(fli1a:Myr-GFP)*^*ncv2Tg*^ (Fukuhara et al., 2014), *Tg(dab2:GFP)*^*ncv67Tg*^ (Shin et al., 2019). *Tg(lyve1:mCherry)*^*ncv87Tg*^ and *Tg(14xUAS:3xFLAG-NTR,NLS-mCherry)*^*ncv514Tg*^ were generated in this study.

### Genotyping

For *pdgfrb*^*um148*^ the following primers were used for PCR

pdfgrb Forward 5’-ATGCGCTAAAGGTGAATTGG-3’

pdgfrb Reverse 5’-GCGTCTGCCATAGTTGAACA-3’

The PCR product was digested with Mbo1 restriction enzyme at 37 °C for 1 h. The digested product was run on 2% agarose gel. The wild type fragment is cut resulting in two bands of 200 bp and 300 bp long; *pdgfrb*^um148^ mutant is not cut and the band is 500 bp long; heterozygous *pdgfrb*^um148^ is a combination of three fragments with bands sizes of 200 bp, 300 bp and 500 bp.

### Immunohistochemistry

Immunohistochemistry was performed according to a previously published protocol (Le Guen et al., 2014; Shin et al., 2016) with the following modifications. After acetone treatment embryos were treated with Proteinase K at 10 mg/ml diluted in PBST for 35 mins. Antibodies used were chicken α-GFP (1:400, ab13970 Abcam), rabbit α-DsRed (1:400, Living colors, 632496 Takara Bio), rabbit α-Phospho-p44/42 MAPK (1:250, #4370 Cell Signaling Technology), and α-rabbit IgG-HRP (1:1000, #7074 Cell Signaling Technology). TUNEL staining was performed with *In Situ* Cell Death Detection Kit, Fluorescein (Merck, 11684795910) with the instruction provided by manufacture.

### Image acquisition

Embryos were anesthetized and mounted in 1% low-melting agarose on a 35-mm-diameter glass-base dish (627870 or 627861 Greiner). Confocal images were obtained using a Leica TCS SP8 confocal microscope (Leica Microsystems) equipped with water-immersion 25X (Fluotar VISR, 0.95 NA) objective, water-immersion 40X (HC PL APO CS2, 1.1 NA) objective and glycerol-immersion 63X (HC PL APO CS2, 1.3 NA) objective or FluoView FV1000/FV1200/FV3000 confocal upright microscope (Olympus) equipped with a water-immersion 20x (XLUMPlanFL, 1.0 NA) lens. The 473 nm (for GFP), 559 nm (for mCherry), and 633 nm (for Qdot 655) laser lines in FluoView FV1000/FV1200/FV3000 confocal microscope and the 488 nm (for GFP) and 587 nm (for mCherry) in Leica TCS SP8 confocal microscope were employed, and 488 nm and 651 nm on the Zeiss NLO710, respectively.

### Image analysis

Image quantification was performed using z-stacks in ImageJ 2.0.0 (Schindelin et al., 2012), Olympus Fluoview (FV10-ASW, FV31S-SW), or IMARISx64 9.5.1 software (Bitplane). Total LEC number was counted manually using the overlay of DsRed and GFP channels over five somites in the trunk. Lymphatic vessel area was calculated by rendering the surface using DsRed channel, the non-lymphatic structures were manually removed. Measurements of surface area were exported directly from Imaris (Bitplane).

#### Cell tracking

To quantify the migrating distance the centre of PL nuclei in figure 3G and figure 4D was manually tracked until either cell dying or disappearing from the view in Imaris (Bitplane). The individual cell track was generated by “spot” and “cell track” function and then manually edited if needed. The data was exported to GraphPad for plotting and statistical analysis.

#### Distance measurement

The migrating distance in all figures were measured in three-dimension using spot function in Imaris (Bitplane), see cell tracking. In figure 3B-C, the sprouting front of PL just migrating out of HM was chosen as start point (T0) and the migrating front at the end of the time lapse video was chosen as end point (T1). A direct line was used to connect the dots and the length of line segment is measured. In figure 2H, the perpendicular distance between point T1 and HM was also measured in addition to the measurement above.

### Chemical treatment

To inhibit Cxcr4 signalling, the embryos were treated in 20 µM antagonist ADM3100 (Merk) from 51 hpf to 72 hpf. To block phosphorylation and activation of ERK1/2, embryos were treated in 10 µM SL327 (EMD Millipore) from 51 hpf to 72 hpf. Embryos were anesthetized and mounted in 1% low-melting agarose in a 2-well slide with separate chambers which allows. The prepared chemical solution (3 ml) was added on top of the agarose layer in one chamber and the E3 water or DMSO (3 ml) to the other.

### Fluorescence Activated Cell Sorting (FACS) and qPCR analysis

Embryos of *Tg(flt1:YFP);Tg(−5*.*2lyve1b:DsRed2)* and *Tg(flt1:YFP);Tg(−5*.*2lyve1b:DsRed2)* were collected at 3 dpf and screened as described in figure 3D, dissociation was performed as previously described (Kartopawiro et al., 2014). The dissociated cells were sorted using a FACS Aria III (BD Biosciences) into 300μl TRIzol™ LS Reagent (Thermo Fisher). Total RNA was extracted using the Quick-RNA Microprep kit (Cambridge Bioscience) following the manufacturer’s instructions. RNA quality and concentration were determined using 2100 Bioanalyser Instrument (Agilent) together with Bioanalyser High Sensitivity RNA Anlysis Kit (Agilent). 1 ng of RNA template was subjected to cDNA synthesis using SuperScript™ VILO™ cDNA Synthesis Kit (Thermo Fisher). qPCR analysis was performed using the primers in Table S1 on CFX384 Touch Real-Time PCR Detection System (BioRad). Data were analysed using the CFX Maestro Software (BioRad). The geometric average of *rpl13* and *β-actin* expression was used as reference to calculate relative gene expression of target genes with the ddCT method. Primer sequences listed below in Table 1.

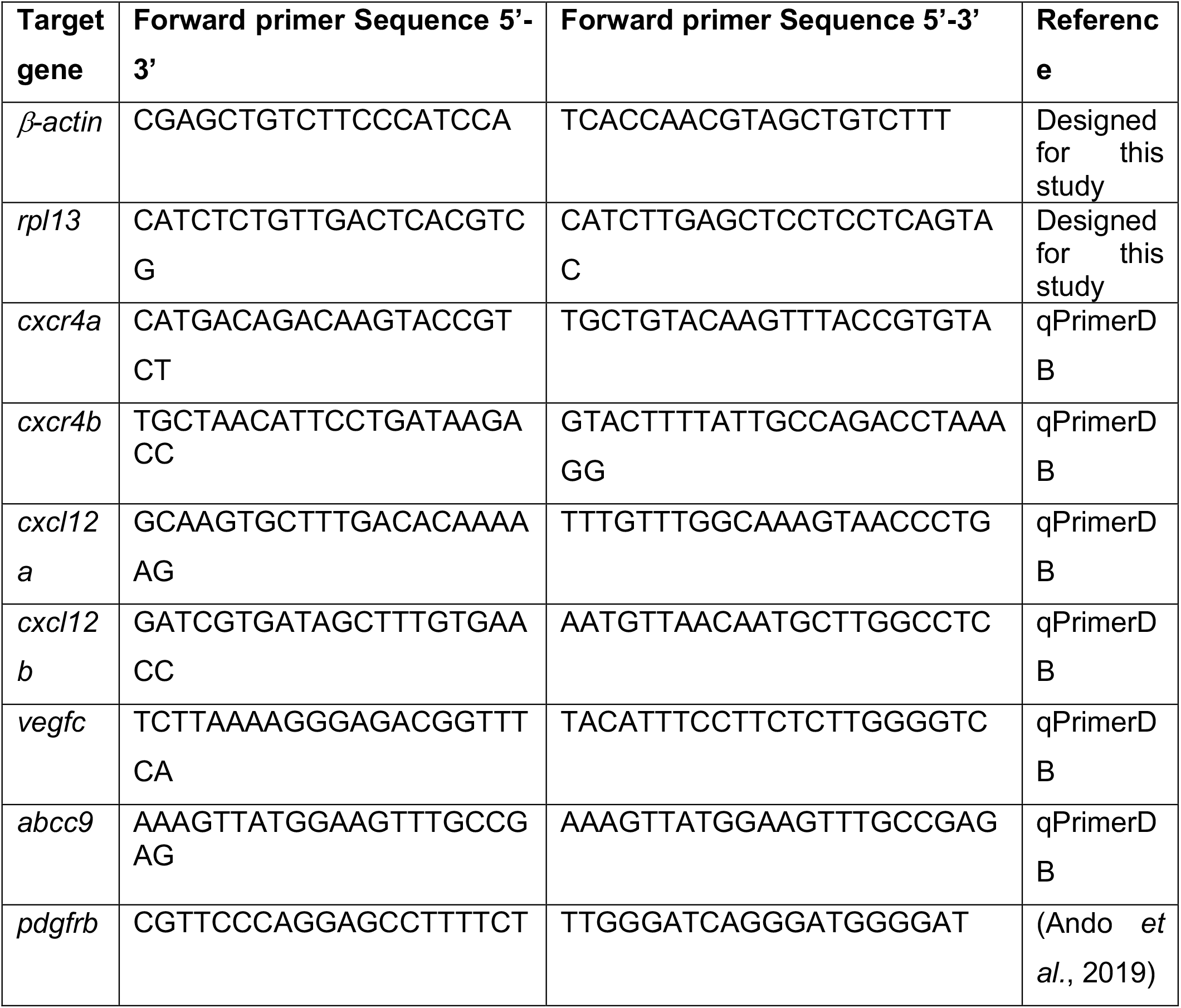

### Ablation experiments

#### Ablation with multi-photon microscopy

For mural cells ablation, embryos of *Tg(−5*.*2lyve1b:DsRed2);Tg(pdgfrb:GFP)* were laterally mounted in 1% low melting-point agarose at 57 hpf. An aISV with migrating LEC was chosen randomly, a GFP-positive MC were identified using 488-nm laser. MC locating ahead of migrating route were ablated using a two-photon laser at 790 nm (Mai Tai, Spectr-Physics Millenia PRO). Control ablations were performed as above but the adjacent area to the pdgfrb+ cell targeted with the two-photon laser. For aISV ablation, *Tg(−5*.*2lyve1b:DsRed2);Tg(flt:YFP);Tg(pdgfrb:GFP)* embryos were prepared as described above and an aISV, with LEC migrating along, was ablated using the two-photon laser targeting the nuclei of aEC. Larvae were imaged before and after ablation with a Zeiss LSM 710 FCS confocal microscope, which was followed by either time-lapse imaging for around 5 h or follow-up z-stack imaging at 3 dpf.

#### Ablation with NTR-Mtz system

Embryos from *Tg(Pdgfrb:Gal4FF,UAS:NTR-mCherry);Tg(fli1a:GFP)* were collected and screened as described in figure 2A. Embryos both positive and negative for *mCherry* were treated with medium mixture of E3 water and either 5mM MTZ or DMSO from 48 hpf, and replaced daily with fresh medium mixture. The region above the yolk extension was imaged at 120 hpf. The formation of thoracic duct was analyzed by scoring the extend of the thoracic duct formation.

### Statistical analysis

Statistical analysis was performed using Prism software (GraphPad). Gaussian distribution of samples was tested with either D’Agostino–Pearson test or Shapiro-Wilk normality test. Student’s t test was used for comparison of two means. One-way ANOVA with post hoc test was used for multiple comparison as stated in corresponding figure legend.

## Acknowledgment

This work was supported by Wallenberg Academy Fellowship (2017.0144), Ragnar Söderbergs Fellowship (M13/17), Vetenskapsådet (VR-MH-2016-01437) and Jeanssons Foundation. The SciLifeLab Zebrafish facility in Uppsala hosted zebrafish. FACS was performed at BioVis at Uppsala University. We are grateful to A. Chiba, H. Nakajima, S. Yuge, T. Babazono, W. Koeda, K. Hiratomi, M. Sone, E. Nakamura, K. Kato, and H. Ichimiya for technical assistance.

## Author contribution

D.P, K.A. and K.K, conceptualised the project, performed, analysed experiments and co-wrote the manuscript. R.S. and M.G. performed and analysed experiments. N. M, C. B., and S.F unpublished reagents and resources contribution.

## Figure Legends

**Supplementary Figure 1.**
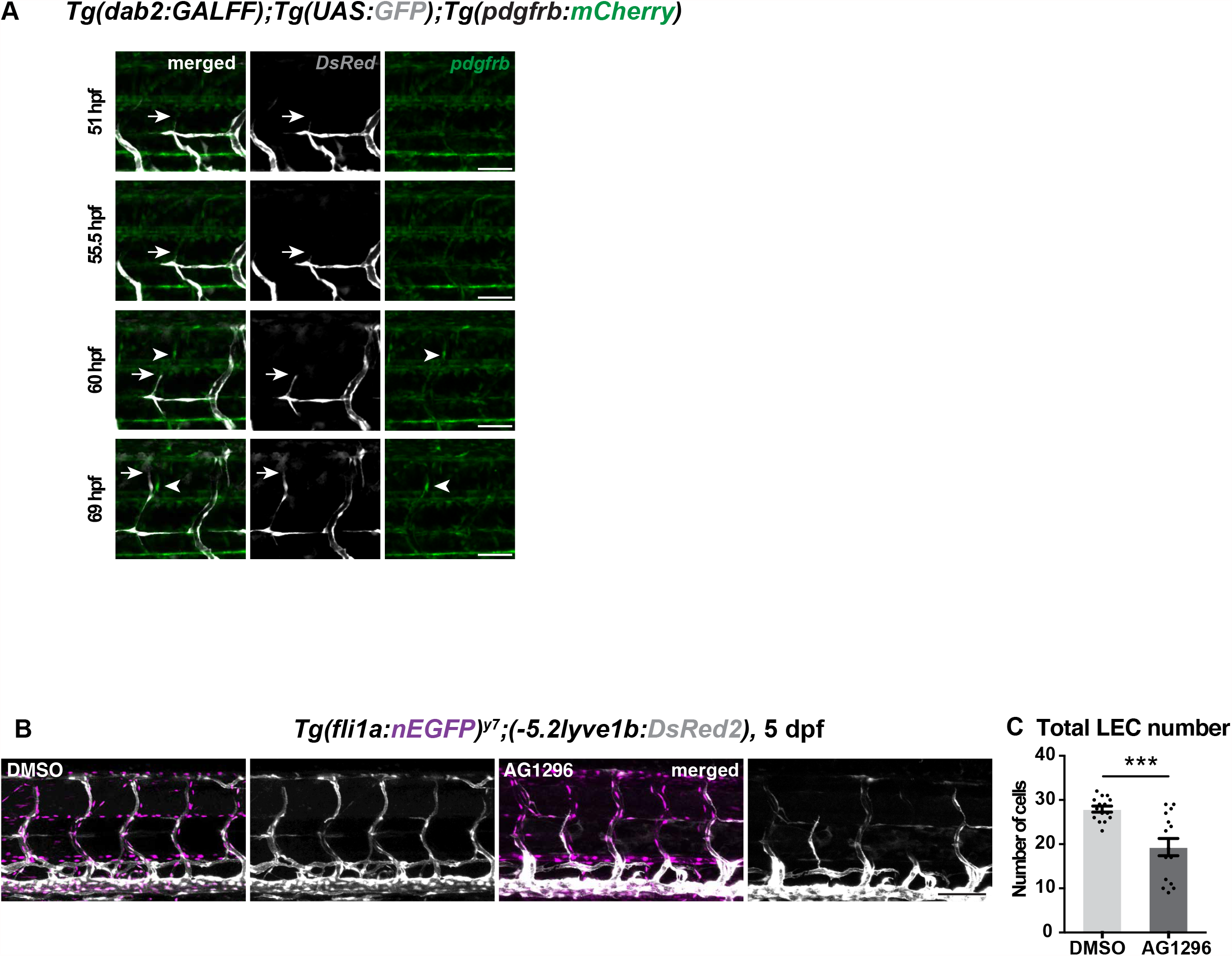
related to Figure 1; *pdgfrb* ^high^ mural cells emerge around aISVs prior to LEC migration and provide guidance. (A) Confocal stack images from time-lapse images in the trunk of *Tg(dab2:GALFF);Tg(UAS:GFP)* in grey and *Tg(pdgfrb:mCherry)* in green. White Arrow heads, MCs appear around aISV. Arrow, sprouting front of migrating LEC. Scale bar; 50 μm. (B) Confocal stack images of 5 dpf *Tg(fli1a:nEGFP);(−5*.*2lyve1b:DsRed2)* treated with 20 μM PDGFR inhibitor AG1296 (n=15) or DMSO (n=15) from 48 hpf. Data are presented as mean ± SEM, unpaired two-tailed Student’s t-test was used. ***p<0.0005.

**Supplementary Figure 2.**
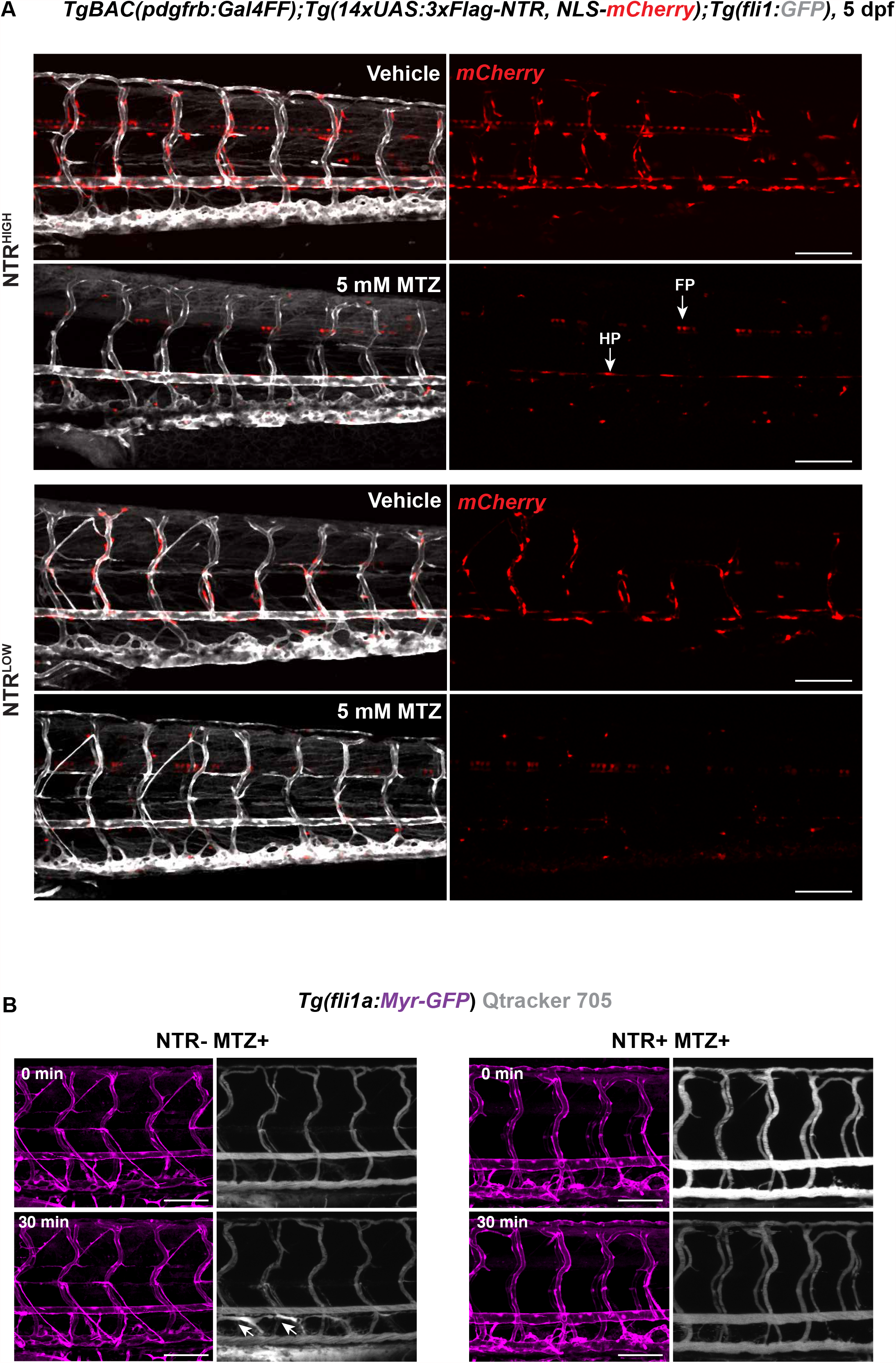
related to Figure 2; Mural cells are required for robust formation of the lymphatic vascular bed. (A) Representative confocal stack images of 5 dpf *TgBAC(pdgfrb:Gal4FF);Tg(14xUAS:3xFlag-NTR, NLS-mCherry);Tg(fli1:GFP)* with high and low NTR expression treated with 5 mM MTZ or vehicle for 16 h. NTR expression was highly selective on MCs but not on *pdgfrb*-low population. MCs were ablated after 16 h of 5 mM MTZ treatment. Arrows indicate floorplate and hyperchord were not ablated by the treatment despite the expression of NTR. Scale bar; 100 μm (B) Confocal stack images from *Tg(fli1a:Myr-GFP)* in control and ablated embryos described in Figure 2A post injection of Qtracker 705 vascular labels (shown in grey) into common cardinal vein. Arrows indicate lymphatic vessels labeled by leaked dye when injected or during the circulation in the control but not in the MC-ablated larva. Scale bar; 100 μm

**Supplementary Figure 3.**
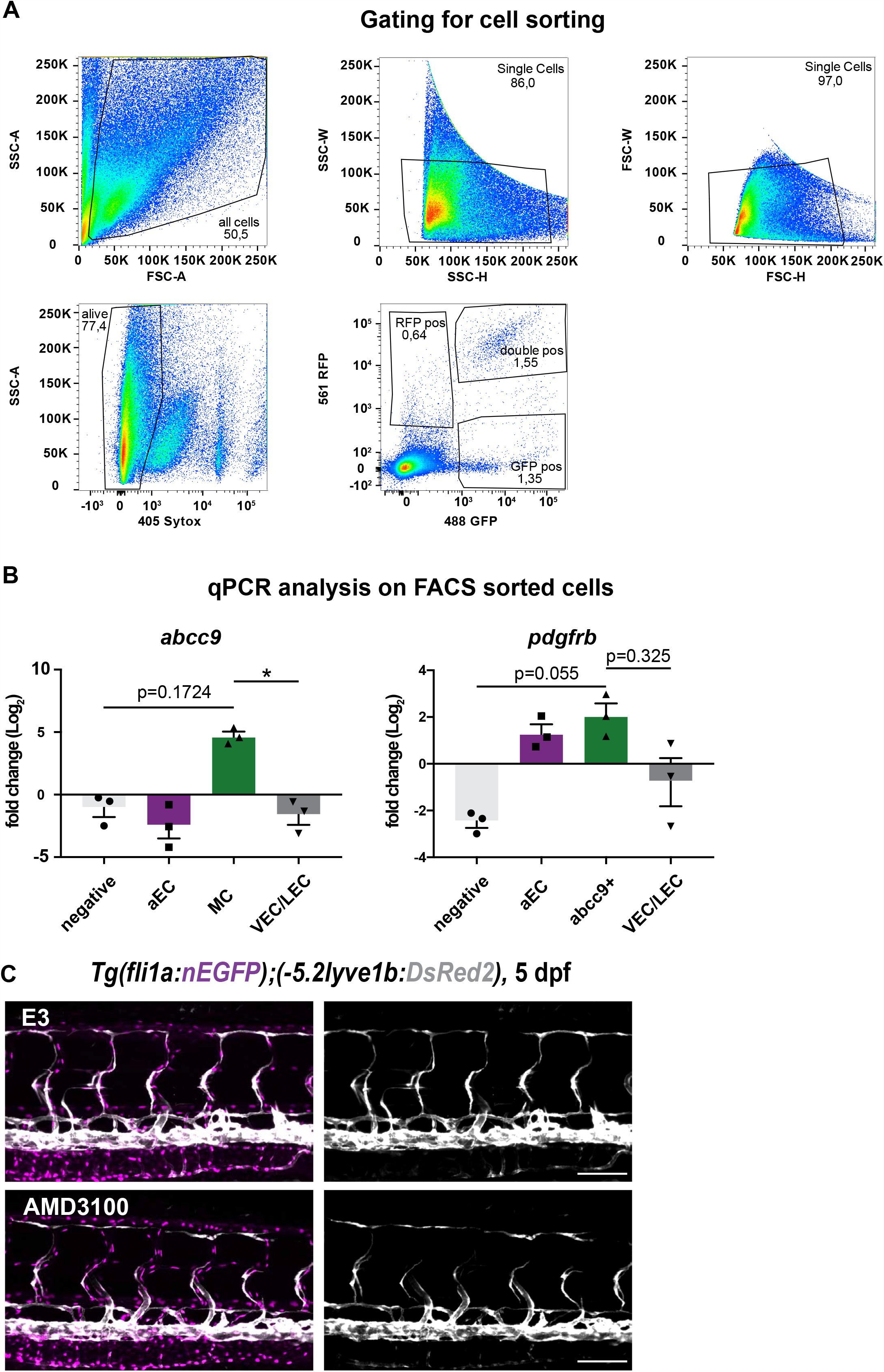
related to Figure 3; Chemokines are expressed in ISV-mural cells during LEC migration. (A) Gating strategy for FACS sort of *Tg(flt1:YFP);Tg(−5*.*2lyve1b:DsRed2) and Tg(abcc9:GFP); Tg(kdrl:DsRed2)* as described in Figure 3D. Sorting was performed on all singlet, alive cells according to their expression of DsRed (red, 561 nm) and GFP/ YFP (green, 488 nm). (B) RT-qPCR of *abcc9* and *pdgfrb* expression in trunk ECs and MCs cells at 3 dpf as described in (3E). Graph represents gene expression relative to gematric average of *rpl13* and *β-actin* from three biological repeats (mean ± SEM). Kruskal-Wallis test with Dunn’s post hoc test was used. No significance (ns), p≥ 0.9999. (C) Confocal stack images of embryos as described in Figure 3F treated with 20 μM ADM3100 or E3 water from 51 hpf to 120 hpf. Scale bar; 100 μm.

**Supplementary Figure 4.**
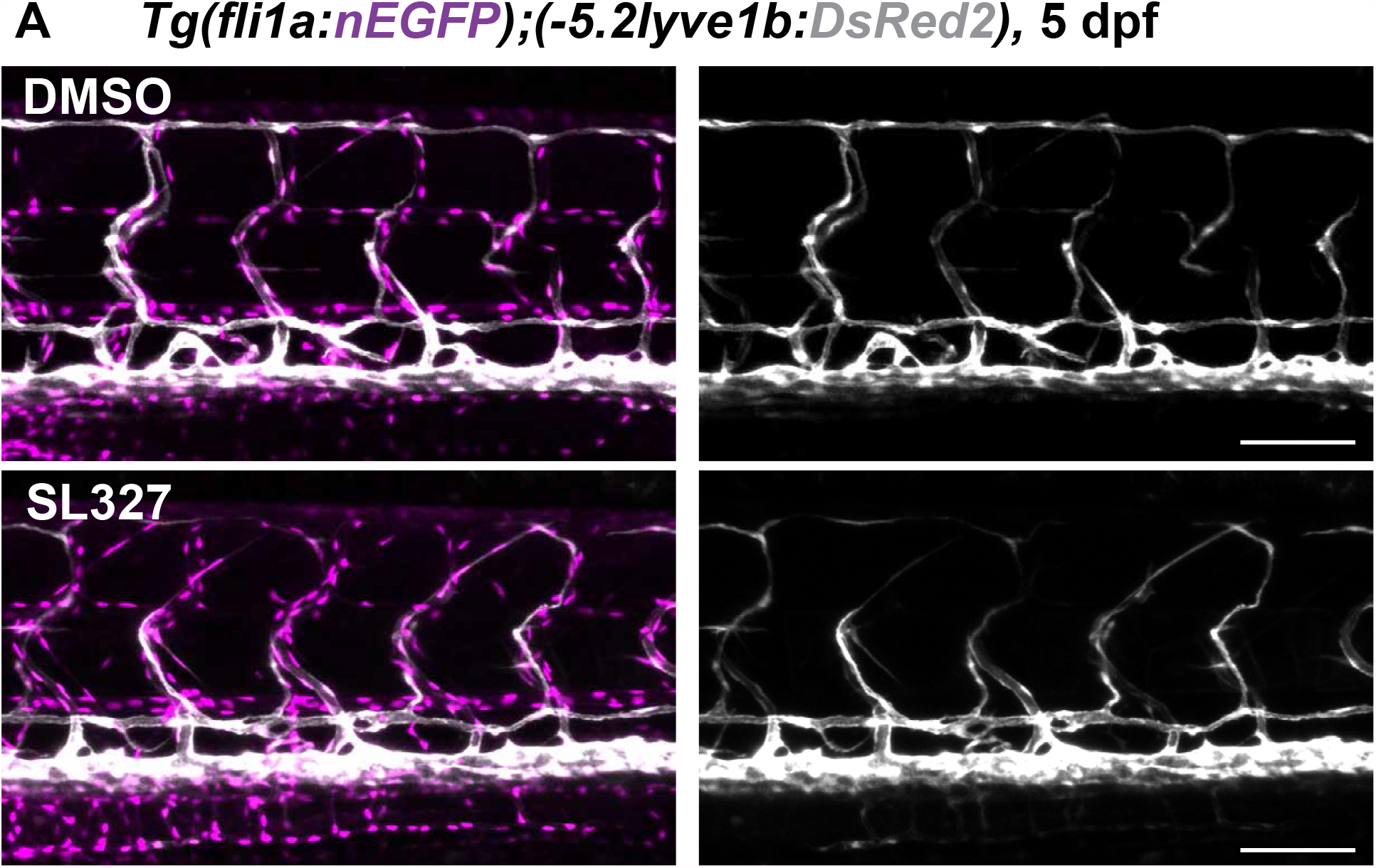
related to Figure 4; Mural cell and arterial derived *vegfc* promotes lymphatic endothelial cell migration and survival. (A) Confocal stack images of embryos as described in Figure 4C treated with 4 μM SL327 or DMSO from 51 hpf to 120 hpf. Scale bar; 100 μm.

**Supplementary Movies S1 related to Figure 1**.

Confocal time lapse imaging in trunk in 2 dpf *Tg(lyve1b: mCherry), Tg(kdrl:TagBFP), TgBAC(pdgfrb:GFP)* embryos corresponding in Figure 1B

**Supplementary Movies S2 related to Figure 1**.

Confocal time lapse imaging in trunk in 2 dpf *TgBAC(pdgfrb:GAL4FF);(UAS:GFP), Tg(−5*.*2lyve1b:DsRed2)* embryos corresponding in Figure 1E

**Supplementary Movies S3-4 related to Figure 1**.

Representative confocal time lapse imaging of LEC migrating with or without contacting MCs in trunk in 2 dpf embryos corresponding Figure 1G.

**Supplementary Movies S5-6 related to Figure 2**.

Representative confocal time lapse imaging of multi-photon ablation of MCs in *TgBAC(pdgfrb:GAL4FF);(UAS:GFP), Tg(−5*.*2lyve1b:DsRed2)*, corresponding Figure 2F.

**Supplementary Movies S7 related to Figure 3**.

Representative confocal time lapse imaging of multi-photon ablation of aISV in *Tg(flt1:YFP)*; *TgBAC(pdgfrb:GFP); Tg(−5*.*2lyve1b:DsRed2)*, corresponding Figure 3A.

**Supplementary Movies S8-11 related to Figure 3**.

Representative confocal time lapse imaging of *Tg(fli1:GFP);Tg(lyve1b:mCherry)* treated with 20 μM ADM3100 or E3 water at from 51 hpf, corresponding Figure 3G.

**Supplementary Movies S12-15 related to Figure 3**.

Representative confocal time lapse imaging of *Tg(fli1a:nEGFP)*^*7y*^; *Tg(−5*.*2lyve1b:DsRed2)* treated with 10 μM SL327 or DMSO from 51 hpf. Grey arrowheads indicate cell death.

## Notes

### Competing Interest Statement

The authors have declared no competing interest.

